# A bug’s tale: revealing the history, biogeography and ecological patterns of 500 years of insect introductions

**DOI:** 10.1101/2022.04.13.488245

**Authors:** Daniela N. López, Eduardo Fuentes-Contreras, Cecilia Ruiz, Sandra Ide, Sergio A. Estay

## Abstract

The arrival of Europeans to the Americas triggered a massive exchange of organisms on a continental scale. This exchange was accelerated by the amazing increase in the movement of people and goods during the 20th century. In Chile, scientific and technical literature contains hundreds of records of exotic insect species in different parts of the territory, from the hyperarid Atacama Desert to the Magallanes Region. Here, we analyze the biological patterns obtained from the database of introductions of exotic insects to Chile from the European arrival to the present. Our task includes a review of old records in museums, libraries, collections, expedition records and catalogs. Almost 600 species of exotic insects have been reported in Chile. Introductions started at the very arrival of Europeans to the central valley of Chile and underwent a huge acceleration in the second half of the 20th century. The order Hemiptera was the most prevalent among introduced insects. Most species are linked to agriculture and forestry. Species are of Palearctic origin in more than 50% of the records. In terms of temporal trends, the rate of introduced species shows an abrupt increase at the beginning of the 1950s. This change may be associated with the strong development in agriculture and forestry in Chile after World War II and the increase in intercontinental air traffic. We believe that the understanding of past patterns of introductions is an important component in the design of current policies to minimize the impact of invasive insects.

## Introduction

The advance of civilizations, human migration and the consolidation of trade between different regions have led to a strong increase in species movement (Buckland 1981). This exchange was accelerated by the amazing increase in the movement of people and goods during the 20th century (Wainhouse 2005). Indeed, the exchange of different species of insects has intensified in the last 200 years, together with the great transcontinental movements of people, goods and services (Mack et al. 2000; Chapman et al. 2017; Liebhold et al. 2017). Today, all countries show hundreds or thousands of alien species established in their ecosystems (Mack 2003; Langor & Sweeney 2009); however, in many cases it is difficult to determine the origin, path and date of the introduction. Furthermore, recent estimates indicate that observational bias means that many introduced pest species still go unreported (Bebber et al. 2019).

Some authors have proposed that social and economic factors are key components in the increase in propagule pressure or species introductions (Santini et al. 2013; Bacon et al. 2014), whereas ecology and biogeography are the main determinants of establishment (Santini et al. 2013; Schulz et al. 2019). In the first case, accelerated economic growth, the agricultural or ornamental use of exotic plants, connectivity (e.g., the number and availability of ports) and access to areas with invasive species emerge intuitively as important variables to explain the rate of insect species arriving in a new country (Dehnen-Schmutz et al. 2007; Hulme 2009; Banks et al. 2015; Chapman et al. 2017; Seebens et al. 2018; Bebber et al. 2019). On the other hand, biogeographic similarity (including climatic similarity), host availability, and community invasiveness have been described as the most important factors to explain the establishment of new insect species (Shea & Chesson 2002; Bacon et al. 2014; Burns 2015; Schulz et al. 2019). Assuming these ideas, it is the particular combination of these factors that will define the configuration of the exotic fauna in a region. For example, the pool of exotic insects in a country could be dominated by species whose origins are in biogeographic regions with environmental conditions that match those of the new habitats. An alternative scenario is a pool of exotic insects that arrived using paths mainly associated with the main economic activities of the country, regardless of their biogeographic origin.

In a similar vein, it is not only the species’ identities that can reflect different processes structuring the exotic community. Diversity of orders or families may or may not respond to ecological factors. Exotic insect communities can be a reflection of the global richness of these groups, with the representation of orders or families being proportional to their world richness, or the exotic richness may be biased toward a group due to socioeconomic or ecological variables (Sailer 1978, 1983; Yamanaka et al. 2015, Bebber et al. 2019).

In the Americas, the arrival of Europeans triggered a massive exchange of organisms on a continental scale. Accidental and intentional introductions of plants and animals promoted the establishment of new insect species, especially those associated with crops of foreign origin. In continental Chile (excluding oceanic islands), new food and crops were introduced in the 16th century (Prado 1991); however, few exotic insects were reportedly established in the country until the 19th century (Prado 1991; Artigas 1995; González 2012). Other routes of introduction of insects were the establishment of ornamental plants, forestry, livestock and accidental transport in human baggage. An important intentional pathway for insect introductions to Chile, especially during the 20th century, was the implementation of biological control programs (González & Rojas 1966; Zúñiga 1985; Rojas 2005). The introduction of these beneficial insects was a response to the increasing impact that exotic pests had on food production.

In Chile, scientific and technical literature contains hundreds of records of exotic insect species in different parts of the territory, from the hyperarid Atacama Desert to the Magallanes Region. Using this information, here we analyze the biological patterns of introductions of exotic insects to Chile from the first European arrival to the present. In particular, we analyze temporal trends, taxonomic diversity, biogeographic origin, and main impacts of these species on different areas of human activity. Through the description of these patterns, we can obtain a better understanding of the process involved in biological invasions over a time scale of centuries.

## Material and methods

### Database

We reviewed records of insect introductions to continental Chile in scientific articles, museums, libraries, collections, expedition records and catalogs. First, we searched for explicit statements of first records of exotic insects in the country. In some cases, we included the first mention of the species in Chile when the authors explicitly recognized that the specific year of the introduction is unknown. We included species that were eradicated by governmental initiatives, but that were originally successfully established in the country. We collected species name, taxonomic position, and year of introduction (or first mention). After we obtained our list, we reviewed the literature to determine origin, type of impact, and if the species was used for biological control. For origin we used the classification of biogeographic Realms from Olson et al. (2001). Given that a species can belong to more than one Realm, we considered all Realms including the native distribution according to its importance. For type of impact on human activities we used the following categories: agriculture, forestry, ornamental, environmental (impacts on biodiversity, endangered or endemic species or ecosystems), livestock, human health and infrastructure (damage to artificial structures, roads, ports, heritage buildings, etc.). To classify the species in one or more categories, we searched for impact descriptions in the scientific and technical literature. In some cases, when no description of impact was found, we recorded the impact as unknown.

### Methods

First, we estimated several descriptive statistics. We calculated the taxonomic distribution of orders and families of the introduced insects, and also the distribution of orders as a function of the dominant Realm of origin. We compared the proportional distribution of orders of introduced insects with the proportional distributions of insect orders in the world and Chile. For world data of insect richness, we used Stork (2018); for Chilean insect richness, we used CONAMA (2008). In the same vein, we estimated the distribution of type of impact. The type of impact acts as a descriptor of the pathway of introduction.

We examined temporal trends in insect introductions. The accumulated number of species was calculated for the complete dataset, main orders, and for those insects considered biological control species (despite an intentional or accidental introduction). For the complete dataset we evaluated three hypotheses of the temporal evolution in species accumulation (S). We compared linear, exponential and segmented trends. A linear accumulation suggests that the rate of accumulation is constant through time (t), independent of changes in population movement, economic growth or/and market changes (no acceleration). Exponential accumulation indicates that species accumulation shows a smooth acceleration (S = a*exp(b*t)), where S is the number of species, t is time and a and b are parameters to be estimated. Finally, a segmented trend points out to an abrupt change in the acceleration of species accumulation, which may be the result of abrupt changes in population movement, economic growth or/and market changes, among other factors. To avoid the noise due to few records in the first centuries, we started our analysis with the number of species accumulated to 1850. We tested these hypotheses by fitting each model to our dataset and selecting the best one using the Bayesian Information Criterion (BIC) in the R environment (R Core Team 2022).

## Results

Our review identified 592 insect species introduced to Chile. Three of them were eradicated after establishment (see SI-1). From this total, we found the approximate date of introduction for 572 species. The exotic insect fauna of Chile is dominated by the order Hemiptera, with almost 40% of the species (Fig. 1a). Coleoptera and Hymenoptera each represent ∼20% of the total species (Fig. 1a). Among the families of Hemiptera, Aphididae and Diaspididae were the most abundant. In Coleoptera, Curculionidae was the dominant family. The distribution of families in Hymenoptera was more homogeneous, with most species acting as a biological control of insect pests. The origin of the introduced species was strongly biased to Palearctic insects. In a secondary position, and well behind Palearctic origin, Nearctic and Neotropical species made important contributions to the exotic insect fauna of Chile (Fig. 1b). For Palearctic, Australasian, Indomalayan and Nearctic species, the dominant order was Hemiptera (Fig. 1b). Only for Neotropical and Afrotropical species was the dominant order Coleoptera (Fig. 1b).

**Figure 1:**
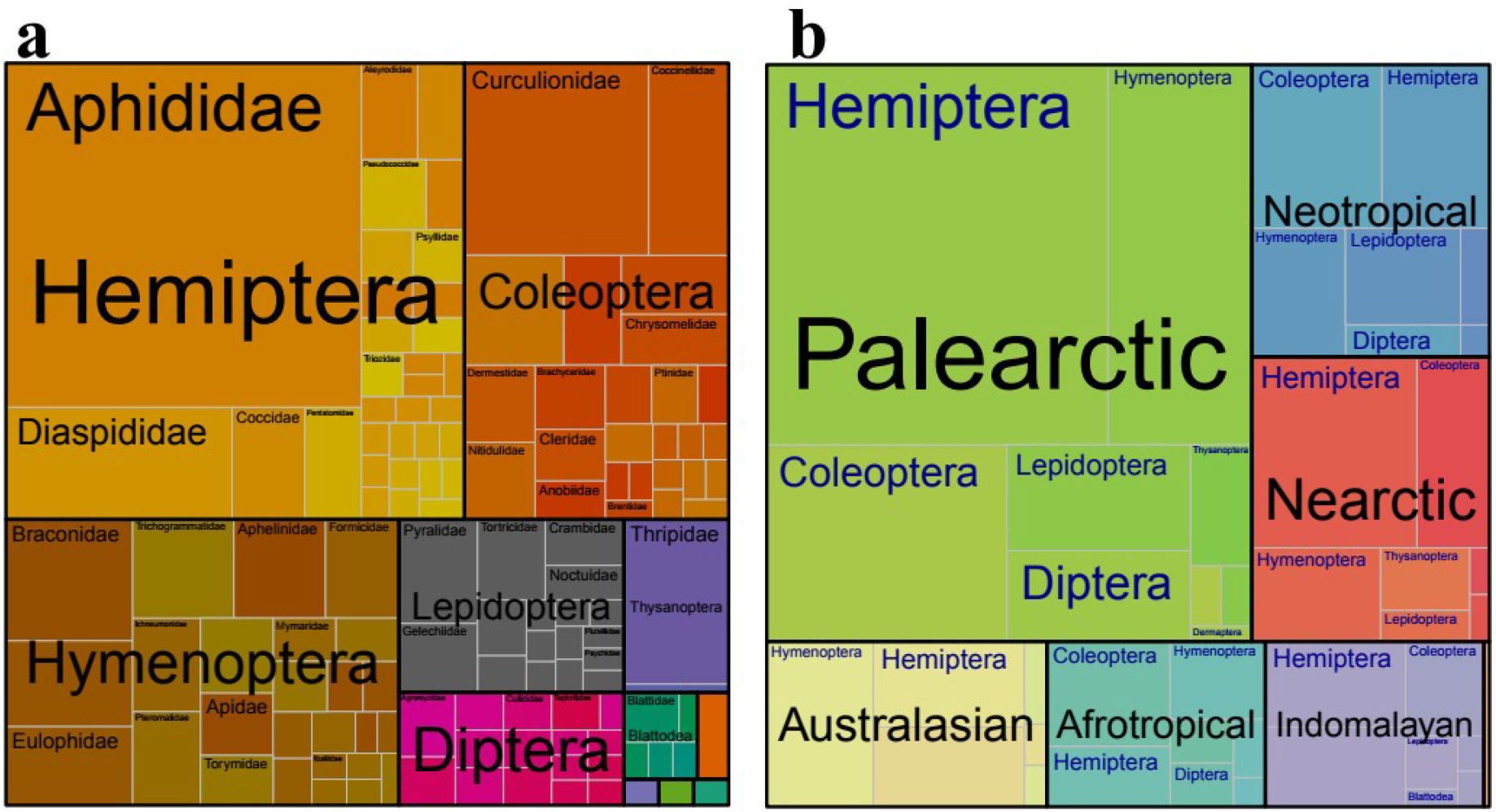
a) Proportional representation of species belonging to orders and families in the exotic insect fauna of Chile. b) Proportional representation of the biogeographic Realms of origin of orders and families of the exotic insect fauna of Chile.

When comparing the relative richness of the introduced insects and world richness, we observe a disproportionate representation of orders Hemiptera and Hymenoptera, and a strong underrepresentation of the orders Coleoptera, Diptera and Lepidoptera (Fig. 2). The same situation occurs when comparing with Chilean insect richness (Fig. 3). Most introduced insects are associated with agriculture, forestry and ornamental areas of impact on human activity (Fig. 4). Other areas show a minor representation.

**Figure 2:**
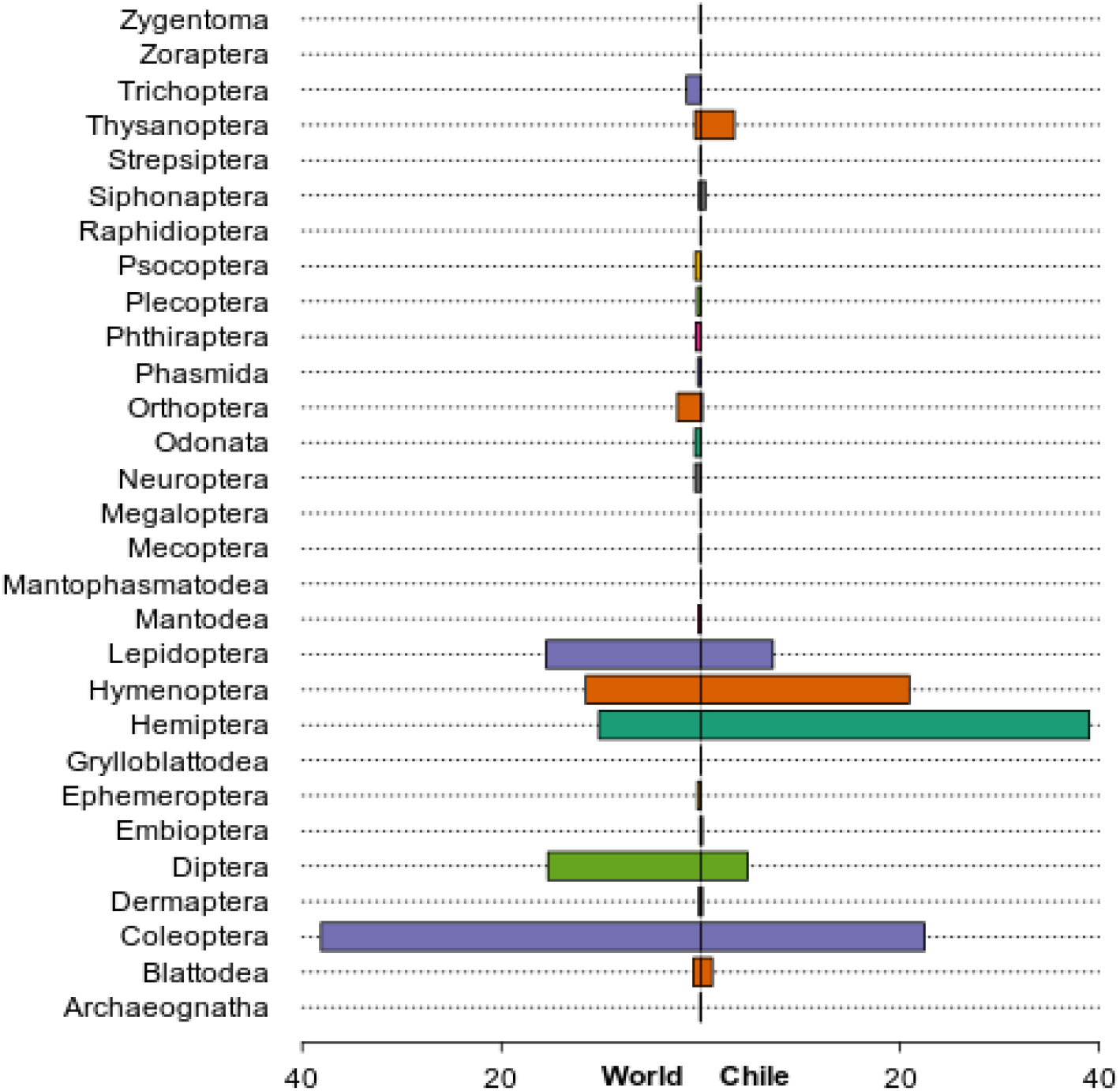
Comparison of the relative species richness of insect orders in the world fauna and in the pool of exotic insects established in Chile.

**Figure 3:**
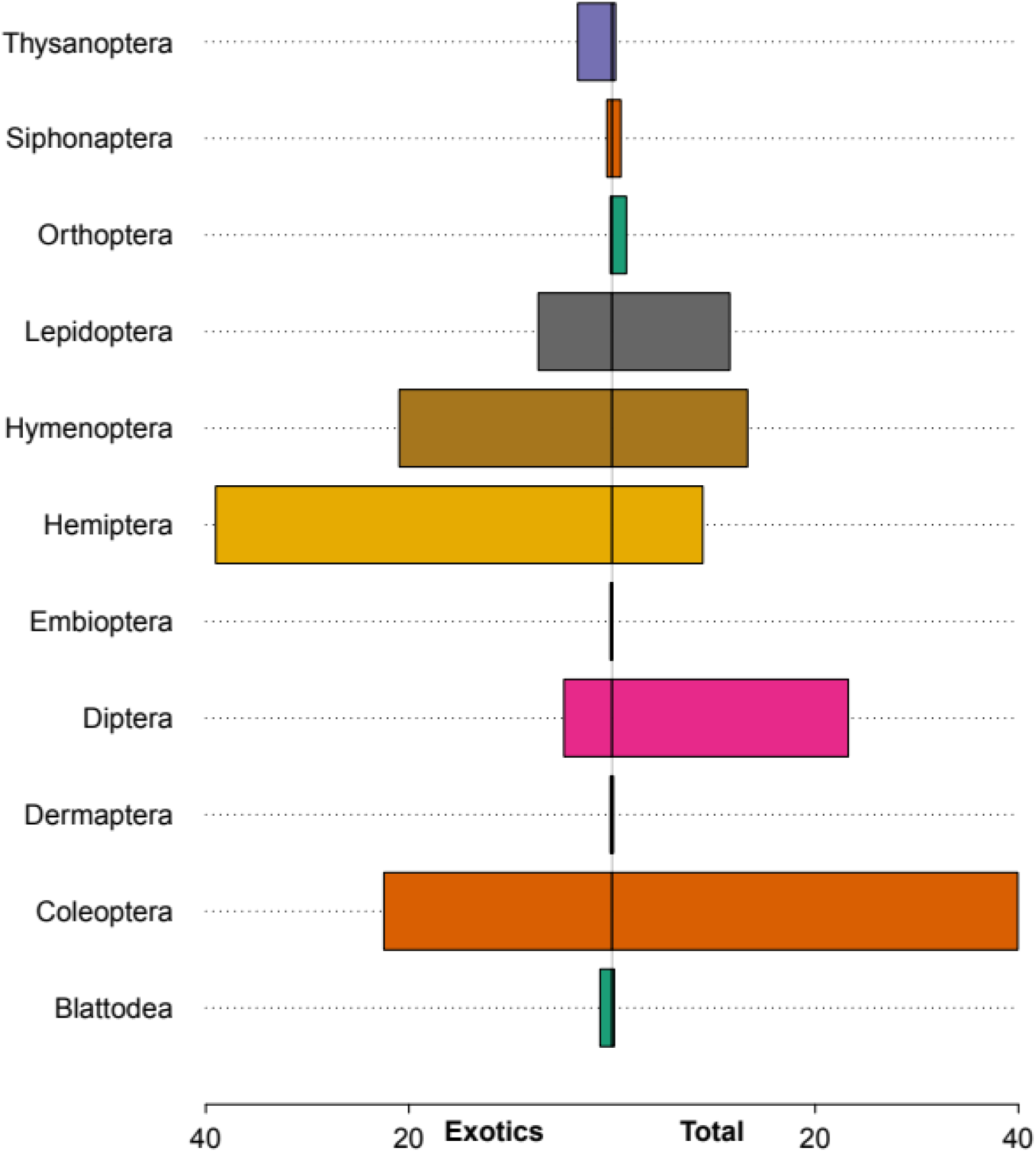
Comparison of the relative species richness of insect orders in Chilean fauna and in the pool of exotic insects established in Chile.

**Figure 4:**
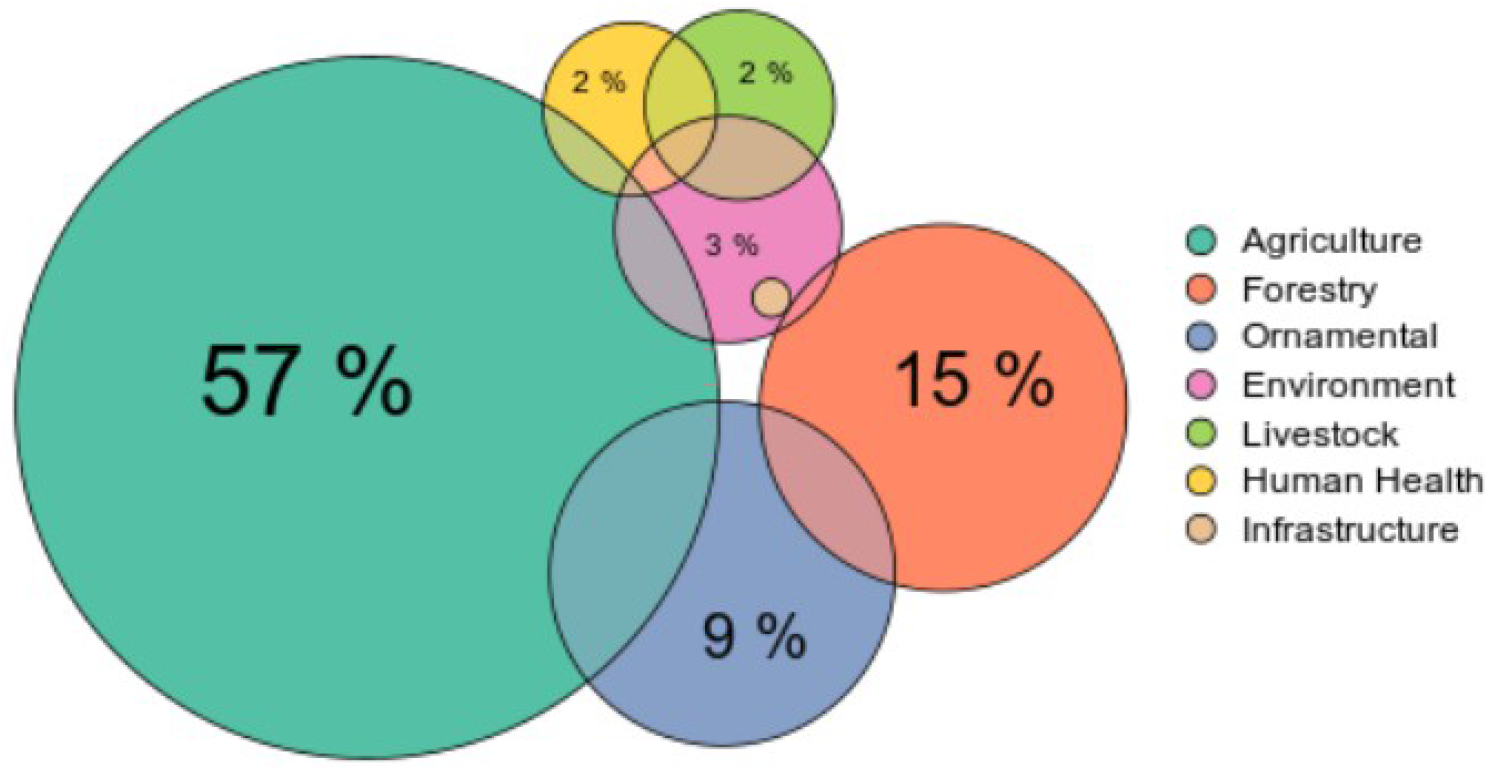
Percentage of species according to the type of impact of exotic insects in Chile. Overlap areas correspond to the percentage of species with more than one area of impact.

When we analyze temporal trends, the best model was the segmented regression (Table 1). The best model showed an abrupt increase in the rate of introduced species around 1949. Starting in that year, the species accumulation rate increased more than three times (Fig. 5). When we look at the more abundant orders, we see that, despite Hemiptera always being the dominant order and showing an accelerated rate of introductions, species accumulation of Coleoptera and Lepidoptera shows a quasi-linear species accumulation (Fig. 6a). Hymenoptera has shown an accelerated increase since the 1950s. (Fig. 6b).

**Table 1.** Results for models fitted to accumulated number of exotic insects in Chile.

| Model | Formula | Parameter | R2 | BIC |
| --- | --- | --- | --- | --- |
| Linear | $S = a + b \cdot t$ | $a = -6219.3; b = 3.312$ | 0.90 | 1472.5 |
| Exponential | $S = a \cdot \exp(b \cdot t)$ | $a \approx 0; b = 0.017$ | 0.99 | 1106.3 |
| Segmented | $S_i = a_i + b_i \cdot t$, with $i = 1, 2$ | $a_1 = -2795.6; b_1 = 1.512; a_2 = -11259.2; b_2 = 5.853$ | 0.99 | 1027.9 |

**Figure 5:**
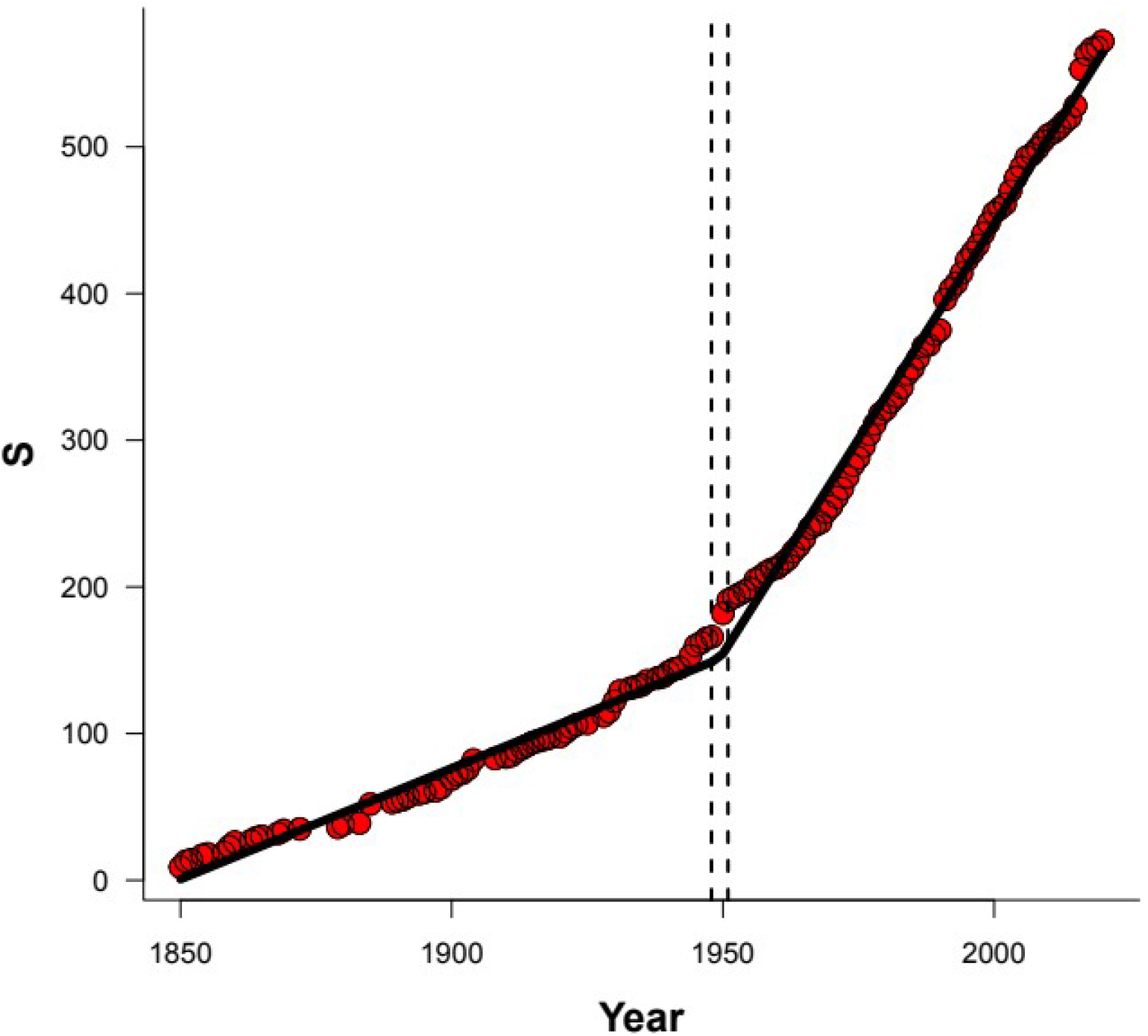
Temporal trend of the accumulated number of exotic species in Chile (S). Red points correspond to observed data. The black line corresponds to the fit of the segmented regression model. Dashed lines correspond to the 95% confidence interval for the break-point in the segmented regression.

**Figure 6:**
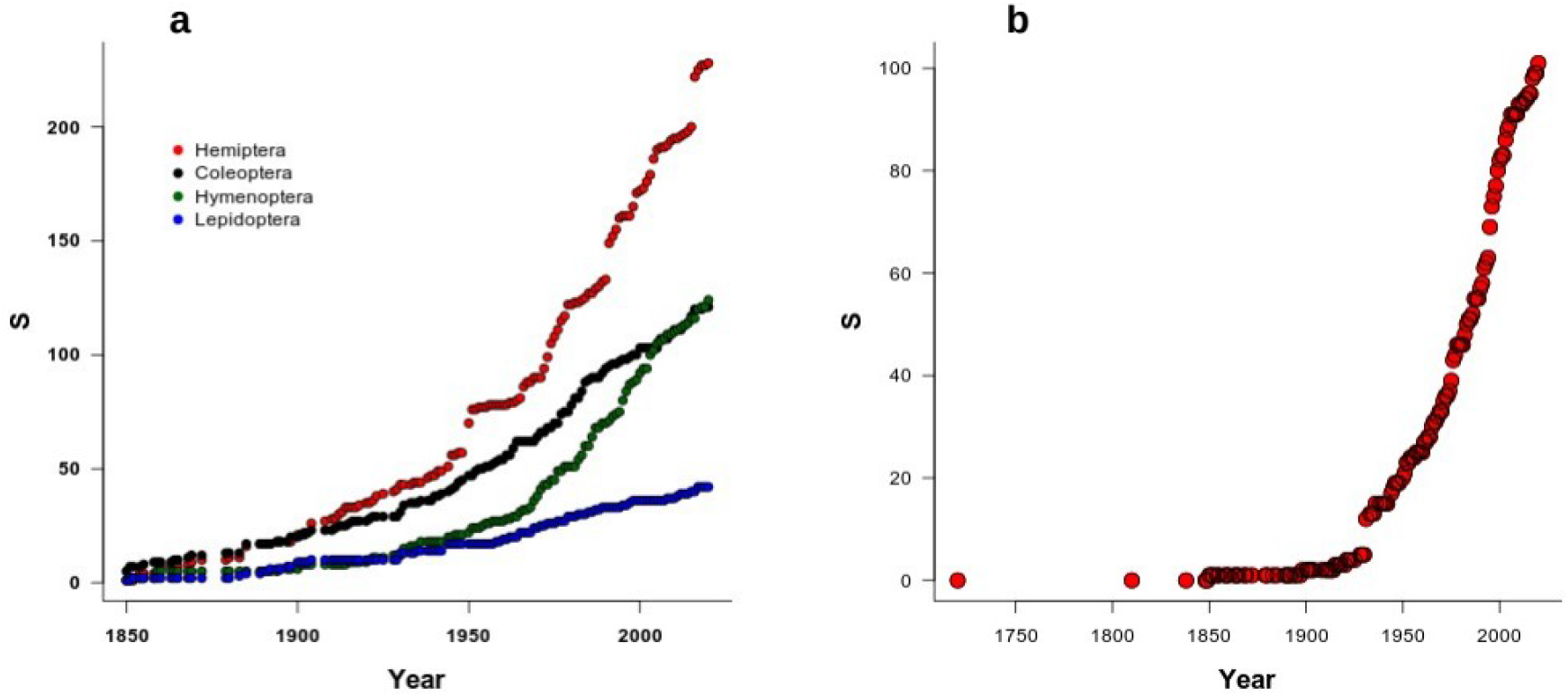
a) Temporal trends in the accumulated number of the four most abundant orders of exotic insects in Chile. b) Temporal trends in the accumulated number of species identified as the biological control in the exotic insect fauna of Chile.

## Discussion

The history of insect introductions to Chile follows the common trend observed all around the world. The arrival of Europeans to the country started major transitions in the insect fauna mainly due to the introduction of exotic plants. For example, the introduction of new crops, such as wheat, began as early as 1541. However, because most exotic plants were introduced as seeds, few insects were detected and very few became a phytosanitary issue before the 19^th^ century (Prado 1991), a similar situation to the one described by Sailer (1983) for the USA. Fruit crops and vines were also introduced early on in colonial times (Lacoste 2004; Lacoste et al. 2011), but major pests of these crops also arrived in Chile during the 19^th^ and 20^th^ centuries (Artigas 1995; González, 2012).

Our results show that the composition of introduced insect fauna in Chile is strongly biased to groups associated with agriculture and forestry. This is a common situation in other parts of the world. Sailer (1978, 1983) and Yamanaka et al. (2015) also showed that most introduced species in North America belong to the order Hemiptera. Waage et al. (2009) also found that Hemiptera (Homoptera in the original) is the order with the most species introduced in Europe and Africa. According to Stork (2018), Hemiptera globally is only the fifth order in terms of the number of species, but it is the most intercepted order in several regions of the world and because its association with crops, forestry and fruit or ornamental trees is disproportionately represented in exotic fauna in many countries (Gippet et al. 2019).

In terms of the origin of the introduced insects, most of them show a Palearctic origin. Species from this Realm represent more than 50% of the introduced insects. Again, this situation has been observed in other parts of the world. Miller et al. (2005) described similar results when they analyzed the richness of exotic scale insects in the USA, all of them associated with agriculture. Yamanaka et al. (2015) also found that in North America and Japan most of the introduced insects are of Palearctic origin. This result is not surprising given that more than 50% of the exotic plants in Chile are of European origin (Fuentes et al. 2014), which supports the hypothesis that the exotic-plant/exotic-insect association is the key promoter of the exotic introduction of arthropods. This is reinforced by the fact that most introduced insects are associated with agricultural, forestry and ornamental plants. Moreover, most early alert systems in the world have been designed to detect exotic insects of economic importance (agriculture, forestry, etc.).

We detect an abrupt increase in the rate of introductions around 1950. Most studies have shown an exponential increase in the rate over years, especially in the last century (e.g., Aukema et al. 2010). In our case, this abrupt change can have several explanations. First, the change in the rate can be a byproduct of the increase in Hemiptera introductions in the same years following the growth in agricultural production post-World War II during the “Green Revolution” (Díaz et al. 2016). A second explanation comes from the major development of biological control programs of plant pests in Chile in the second half of the 20^th^ century, with particular relevance for the parasitoid Hymenoptera (Rojas 2005). Both explanations make reference to changes in agricultural production, but a third alternative is related to the strong increase in air transport (Díaz et al. 2016). The use of international air transport by Chileans showed a marked, strong growth at the beginning of the 1950s. Furthermore, international trade in Chile also increased in the last decades of the 20^th^ century, along with globalization. In particular, such a recent increase in trade with Asian countries could be incorporating new regions with new pools of potential invasive species (Seebens et al. 2018). Finally, observational bias in the report of new introduced species is probably present in our analyses (Bebber et al. 2019). Reports of new introduced species of insects require trained entomologists and international collaboration with specialist taxonomists. For Chile in the 19^th^ century, these human resources were a few foreign naturalists working in the country. The first applied entomologists appeared at the end of 19^th^ and the first half of the 20^th^ century. Finally, a more robust and permanent process of training and networking of applied entomologists and agriculturalist scientists was promoted only during the second half of the 20^th^ century (Artigas 1995; del Pozo et al. 2021). All these variables might be associated with the increase in introduced species, but more detailed analyses are needed to evaluate their relative contribution.

Nowadays, climate change has acted as a promoter of the range expansion of many insect species. For exotic species, ongoing and future climate change could facilitate the short distance dispersion of exotic insects across national borders (Pearson 2006; Hulme 2016). However, for some authors, climate-tracking species should not be considered invasive (Urban 2020).

In this study, we reconstructed the main patterns of insect introductions to Chile. To the best of our knowledge, this is the first attempt to examine the problem of insect introductions from a historical perspective in South America, including all exotic species. This database is the first attempt to compile this information, but this is essentially a work in progress. It has to be updated and improved by governmental agencies, academics and specialists for a better understanding of it. We think that some of the results presented in this study may be representative of other countries in South America. Similarities with other regions suggest that the processes behind insect introductions are common around the world, and their detailed description can be a fundamental tool for managing current introductions and preventing major economic, social or environmental damage.

## Acknowledgments

The authors were supported by ANID PIA/BASAL FB0002 and Fondecyt 1211114

